# Caffeine Increases the Reinforcing Efficacy of Alcohol, an Effect that is Independent of Dopamine D_2_ Receptor Function

**DOI:** 10.1101/2020.09.04.283465

**Authors:** Sarah E. Holstein, Gillian A. Barkell, Megan R. Young

## Abstract

The rising popularity of alcohol mixed with energy drinks (AmEDs) has become a significant public health concern, with AmED users reporting higher levels of alcohol intake than non-AmED users. One mechanism proposed to explain heightened levels of alcohol intake in AmED users is that the high levels of caffeine found in energy drinks may increase the reinforcing properties of alcohol, an effect which may be dependent on interactions between adenosine signaling pathways and the dopamine D_2_ receptor. Therefore, the purpose of the current study was to confirm whether caffeine increases the reinforcing efficacy of alcohol using both fixed ratio (FR) and progressive ratio (PR) designs, and to investigate a potential role of the dopamine D_2_ receptor in caffeine’s reinforcement-enhancing effects. Male Long Evans rats were trained to self-administer a sweetened alcohol or sucrose solution on an FR2 schedule of reinforcement. Pretreatment with caffeine (5-10 mg/kg) significantly increased operant responding for the sweetened alcohol reinforcer, but not sucrose. PR tests of motivation for alcohol or sucrose likewise confirmed a caffeine-dependent increase in motivation for a sweetened alcohol solution, but not sucrose. However, the D_2_ receptor antagonist eticlopride did not block the reinforcement-enhancing effects of caffeine using either an FR or PR schedule of reinforcement. Taken together, these results support the hypothesis that caffeine increases the reinforcing efficacy of alcohol, which may explain caffeine-induced increases in alcohol intake. However, the reinforcement-enhancing effects of caffeine appear to be independent of D_2_ receptor function.

## 1. Introduction

Caffeine is one of the most widely abused psychoactive substances, with an estimated 85% of the U.S. population consuming a caffeinated beverage (Daly and Fredholm, 1998; Mitchell et al., 2014; Nehlig, 1999). However, it is the rapidly rising popularity of highly caffeinated energy drinks (EDs), and their use with alcohol, that has caused concern (De Sanctis et al., 2017; Marczinski and Fillmore, 2014; Vida and Rácz, 2015). Among college-aged adults, estimates of AmED (alcohol mixed with an ED) use range from 8 - 65% (Verster et al., 2018), with approximately one-quarter of alcohol users consuming an AmED in the past month (Brache and Stockwell, 2011; O’Brien et al., 2008). Of particular concern, AmED users reported drinking more alcohol per drinking occasion and drinking longer per drinking occasion than non-AmED users, and experienced more adverse consequences of alcohol use than non-AmED users (Brache and Stockwell, 2011; McKetin et al., 2015; O’Brien et al., 2008; Peacock et al., 2012; Price et al., 2010; Thombs et al., 2010).

Recently, researchers have proposed that individuals may mix alcohol with a caffeinated ED because they find alcohol more pleasurable when combined with caffeine (Marczinski, 2014, 2015; Marczinski et al., 2016). In support of this hypothesis, participants often cite hedonistic motives for combining alcohol with a caffeinated ED, such as reporting AmED use to “increase the pleasure of intoxication”, as well as liking the effect of caffeine and alcohol combined (Droste et al., 2014; Peacock et al., 2015, 2013). Participants are also more likely to choose a caffeinated alcoholic beverage over a non-caffeinated one if given the choice (Sweeney et al., 2017), and both caffeine and caffeinated EDs increase subjective ratings of “liking” for alcohol and increase participants’ desire for alcohol (Heinz et al., 2013; Marczinski et al., 2016; McKetin and Coen, 2014). As caffeine and EDs appear to increase the subjective pleasurable effects of alcohol, as well as motivation for an alcoholic beverage, these results provide preliminary evidence supporting a caffeine-induced increase in the reinforcing efficacy of alcohol (Bickel et al., 2000; Marczinski, 2015, 2014; Marczinski et al., 2016; Stafford et al., 1998).

Animal studies have likewise demonstrated a caffeine-induced increase in alcohol drinking, though these effects are dose- and procedure-dependent (Franklin et al., 2013; Fritz et al., 2016; Kunin et al., 2000; Okhuarobo et al., 2018; Rezvani et al., 2013; SanMiguel et al., 2019). However, the majority of these studies have used non-operant drinking paradigms which do not specifically address the positive reinforcing effects of alcohol or the subject’s motivation for alcohol (Sanchis-Segura and Spanagel, 2006; Tabakoff and Hoffman, 2000). Rather, operant self-administration methods, including both fixed ratio (FR) and progressive ratio (PR) designs, are needed to directly address the impact of caffeine on the reinforcing efficacy of alcohol (Bickel et al., 2000; Stafford et al., 1998). Recent evidence supports a caffeine-induced increase in the positive reinforcing effects of alcohol, as rats self-administered alcohol at higher rates when it was combined with a caffeinated ED, as compared to alcohol alone (Roldán et al., 2018). However, caffeinated EDs include a variety of other potentially psychoactive substances that may likewise alter alcohol intake (De Sanctis et al., 2017; Seifert et al., 2011). More research is therefore needed to demonstrate a direct effect of caffeine on the reinforcing efficacy of alcohol.

Caffeine has also been reported to increase the reinforcing efficacy of both drug (Gannon et al., 2019, 2018; Prieto et al., 2016; Rezvani et al., 2013; Schenk et al., 1994; Shoaib et al., 1999) and non-drug (Bradley and Palmatier, 2019; Retzbach et al., 2014; Sheppard et al., 2012) reinforcers, which has led to caffeine being described as a generalized reinforcement enhancer (Garber and Lustig, 2011; Prieto et al., 2016; Sheppard et al., 2012). Although caffeine is a nonspecific adenosine receptor antagonist, targeting both A_1_ and A_2A_ receptors (A_1_R, A_2A_R) in the brain (Daly and Fredholm, 1998; Fredholm et al., 1999), it is the interaction of adenosine and dopamine signaling pathways that has been proposed to underlie this reinforcement-enhancing effect of caffeine (Ferré, 2016, 2013; Ferré and O’Brien, 2011; Garrett and Griffiths, 1997; Green and Schenk, 2002; Marczinski, 2014). A direct action of caffeine on dopamine signaling has been reported, with caffeine eliciting an increase in extracellular dopamine concentrations within the nucleus accumbens (NAc) (Malave and Broderick, 2014; Quarta et al., 2004; Solinas et al., 2002); however, other studies have not been able to replicate this effect (Acquas, 2002; De Luca et al., 2007; Volkow et al., 2015). Caffeine has also been proposed to indirectly increase dopamine signaling via antagonistic adenosine-dopamine receptor interactions. Within the NAc, adenosine A_2A_Rs and dopamine D_2_ receptors (D_2_Rs) are often colocalized in the form of a heterotetrameric complex (Bonaventura et al., 2015; Ferré et al., 2018b, 2018a). A_2A_R antagonists, including caffeine, increase the affinity and efficacy of dopamine at the D_2_R (Bonaventura et al., 2015; Volkow et al., 2015), suggesting an alternative mechanism by which caffeine can modulate dopamine signaling and enhance the reinforcing efficacy of alcohol and other drugs of abuse. In support of this hypothesis, D_2_R antagonists have been shown to block caffeine-induced increases in cocaine seeking (Green and Schenk, 2002), suggesting that caffeine’s reinforcement-enhancing effects are dependent on D_2_R function.

The A_2A_R-D_2_R heteromer is an intriguing candidate mechanism for the caffeine-dependent increase in alcohol drinking, as previous studies have shown that caffeine-induced increases in alcohol intake can be mimicked by low doses of an A_2A_R antagonist (Arolfo et al., 2004; Micioni Di Bonaventura et al., 2012; SanMiguel et al., 2019) and D_2_R function is critical for the positive reinforcing effects of alcohol (Arolfo et al., 2004; Hodge et al., 1997; Risinger et al., 2000). As alcohol self-administration elicits an increase in extracellular dopamine levels in the NAc (Doyon et al., 2005, 2003; Melendez et al., 2002), researchers have proposed that a caffeine-induced inhibition of A_2A_RS and subsequent increase in D_2_R function may potentiate alcohol-induced increases in dopamine signaling, resulting in an increase in the reinforcing efficacy of alcohol and an increase in alcohol intake (Ferré, 2016, 2013; Ferré and O’Brien, 2011). However, to date, this hypothesis has not been directly tested.

Therefore, to address this gap in knowledge, the current series of studies was designed to examine if an acute administration of caffeine would directly increase the reinforcing efficacy of alcohol and whether this reinforcement-enhancing effect of caffeine was dependent on D_2_R function. Reinforcing efficacy is typically evaluated through changes in both response rate (via an FR schedule of reinforcement) and break point (via a PR schedule of reinforcement) (Bickel et al., 2000; Stafford et al., 1998). Therefore, in order to evaluate the effect of caffeine on the reinforcing efficacy of alcohol, caffeine-induced changes in alcohol-reinforced responding and break point were measured using both FR and PR schedules of reinforcement. As caffeine has been found to increase the reinforcing efficacy of multiple drugs of abuse (Gannon et al., 2019, 2018; Prieto et al., 2016; Rezvani et al., 2013; Schenk et al., 1994; Shoaib et al., 1999), and a caffeinated ED increased operant responding for alcohol (Roldán et al., 2018), we hypothesized that caffeine would likewise increase alcohol-reinforced responding and motivation for alcohol. To determine the contribution of dopamine D_2_Rs to caffeine’s reinforcement-enhancing effects, we evaluated the impact of a D_2_R antagonist (eticlopride) on caffeine-induced increases in alcohol-reinforced responding and motivation for alcohol. Given the proposed role of A_2A_R-D_2_R heteromers to caffeine’s reinforcement-enhancing effects (Ferré, 2016, 2013; Ferré and O’Brien, 2011), we hypothesized that eticlopride would block a caffeine-induced increase in responding for alcohol.

## 2. Materials and methods

### 2.1. Subjects

Male Long Evans rats (50-74 g, Envigo, Indianapolis, IN) were group housed (4/cage) in standard polycarbonate cages with shredded aspen shavings (NEPCO, Warrensburg, NY). Food (Laboratory Rodent Diet 5001, LabDiet, St. Louis, MO) and water were available *ad libitum,* except where noted. Animals were handled daily and were pair-housed approximately 1-2 weeks before the beginning of operant training. The animal colony was maintained on a 12:12 h L:D cycle (lights on between 11AM - 12PM) at 21 ± 2°C. Behavioral testing occurred during the light cycle, beginning at least 2 h after lights on. All experiments were approved by the Lycoming College Institutional Animal Care and Use Committee and complied with the National Institutes of Health guidelines for the care and use of laboratory animals (National Research Council, 2011).

### 2.2. Apparatus

Four operant conditioning chambers (Med Associates, Inc., St. Albans, VT), measuring 29.5 x 24.8 x 18.7 cm (*l* x *w* x *h*) were housed in sound-attenuating chambers equipped with a ventilation fan to mask external noises. Consistent with previously published methods (Faccidomo et al., 2009; Makhijani et al., 2018), a liquid receptacle equipped with a fluid cup and head entry detector was located on one wall with a retractable lever (active lever) located to the right or left of the receptacle. A second retractable lever (inactive lever) was located on the opposing wall, directly opposite the active lever. The location of the active lever and liquid receptacle was counterbalanced across subjects. Responses on the active lever resulted in the activation of a syringe pump and delivery of 0.1 ml of the reinforcing solution to the fluid cup over 1.66 sec. A 2.5 cm diameter white stimulus light, located above the active lever, was illuminated while the syringe pump was active. Responses on the inactive lever were recorded but did not result in any programmed consequences.

### 2.3. Drugs

Alcohol (95% w/v ethanol stock, Pharmco Products Inc., Brookfield, CT) and sucrose were diluted in distilled water. Caffeine and the dopamine D_2_R antagonist eticlopride (S-(–)-3-Chloro-5-ethyl-N-[(1-ethyl-2-pyrrolidinyl)methyl]-6-hydroxy-2-methoxybenzamide hydrochloride; Sigma-Aldrich, St. Louis, MO) were dissolved in saline (0.9% NaCl; Fisher Scientific, Pittsburgh, PA). All drug injections were prepared on the day of testing and were administered intraperitoneally (i.p.) at a volume of 1 ml/kg body weight.

### 2.4. Procedure

#### 2.4.1. Operant self-administration training

Operant self-administration training for a 10% w/v sucrose solution (10S) began once rats reached approximately 280-290 g. Subjects were water restricted for 23.5-24 h prior to the first 16-h overnight training session, consistent with previously published methods (Besheer et al., 2008; Cannady et al., 2013). To encourage responding, the fluid cup was baited with 2 free reinforcers. Responses on the active lever were reinforced on an FR1 schedule for the first 10 reinforcers; the schedule of reinforcement was then increased to FR2. The overnight session ended when subjects had earned 480 reinforcers or 16 h had elapsed (whichever occurred first). If a subject had earned at least 250 reinforcers during the overnight session, it was returned to the home cage with food and water available *ad libitum* and sucrose fading began the following day. However, if fewer than 250 reinforcers were earned, the subject was given access to water in the home cage for 1 h and a second 16-h overnight session was conducted. Rats received up to 3 overnight sessions before advancing to sucrose fading.

#### 2.4.2. Sucrose fading and baseline self-administration

Thirty-minute self-administration sessions were conducted daily (Monday-Friday) on an FR2 schedule of reinforcement, with food and water available *ad libitum.* Fluid cups were baited with 1 free reinforcer prior to the start of each self-administration session to encourage reinforcer-seeking behaviors. Fluid cups were inspected following the completion of each session to confirm intake and any remaining fluid was subtracted from the estimated intake for that session. To establish stable responding for alcohol, a modified sucrose fading procedure was used (Besheer et al., 2008; Samson, 1986). Subjects were initially given 4 self-administration sessions at 10S to stabilize operant responding. Alcohol (A) was then gradually added to the reinforcer as sucrose was gradually eliminated, with subjects receiving at least 2 sessions at each concentration (10S, 2A10S, 5A10S, 10A10S, 10A7.5S, 10A5S, 10A2S, 10A, 10A2S). For 2 subjects that showed low responding following sucrose fading (Experiment 1), additional training sessions at 10A5S and 10A3.5S were included to increase responding for the final 10A2S reinforcer. A lightly sweetened alcohol reinforcer (10A2S) was selected as the reinforcing solution for the remainder of the experiment as it resulted in higher operant response rates and moderate levels of alcohol consumption (0.5-0.8 g/kg), as compared to alcohol alone.

Rats trained to self-administer sucrose received a similar sucrose-fading procedure (Cannady et al., 2013), with at least 4 self-administration sessions at 10S to stabilize responding. Subsequently, sucrose concentrations were gradually reduced (10S, 7.5S, 5S, 2S, 1S, 0.8S), with at least 2 sessions at each concentration before remaining at 0.8S for the remainder of the study. This final concentration was chosen to match active lever responses between the sweetened alcohol and sucrose reinforcers (and thereby match reinforcer efficacy). Following sucrose fading, baseline responding for the 10A2S or 0.8S solutions continued for a minimum of 28 sessions prior to the initiation of saline habituation injections and drug testing.

#### 2.4.3. Experiment 1. Effect of acute caffeine administration on operant responding for a sweetened alcohol solution

Eight rats were used for this study. In an effort to reduce animal numbers, rats in Experiment 1 were also used for Experiments 2 and 6. Two rats were single-housed (prior to the start of caffeine testing) due to fighting. Following baseline drinking, subjects were habituated to handling and saline injections once weekly for 3 weeks, with saline injected 30 min prior to the self-administration session. In order to test the effect of acute caffeine administration on alcohol-reinforced responding, caffeine (0, 5, 10, 20 mg/kg) was administered 30 min prior to the operant self-administration session using a within-subjects, Latin Square design. Doses and pretreatment times were selected based on pilot studies and previous research demonstrating a caffeine-induced increase in alcohol and/or sucrose intake within this dose range (Kunin et al., 2000; Retzbach et al., 2014). Caffeine testing occurred once per week.

#### 2.4.4. Experiment 2. Effect of acute caffeine administration on motivation for a sweetened alcohol reinforcer

Two weeks following completion of caffeine testing (Experiment 1), the effect of caffeine on motivation for a 10A2S reinforcer was evaluated using a progressive ratio 1 (PR1) schedule of reinforcement (Besheer et al., 2008). The PR1 procedure was programmed to increase the response requirement by a factor of 1 for every reinforcer delivered (FR1 for the first reinforcer, FR2 for the next reinforcer, and so on) and previous research has shown that this procedure can be successfully used to repeatedly test motivation for a sweetened alcohol reinforcer (Besheer et al., 2008). For this experiment, a fixed 30-min session was used in order to compare caffeine-induced changes in response rate over time. Therefore, break point in this experiment was operationally defined as the highest response requirement completed in the 30-min session. Subjects received once-weekly injections of caffeine (0, 5, or 10 mg/kg) 30 min prior to the PR session, with dose order randomized across subjects. Caffeine doses were selected based upon the results of Experiment 1. On non-injection days rats completed standard 30-min FR2 training sessions to maintain responding between PR test sessions.

#### 2.4.5. Experiment 3. Effect of acute caffeine administration on operant responding for sucrose

To determine whether caffeine’s reinforcement-enhancing effects were selective for alcohol, a behavior-matched cohort of Long-Evans rats (n = 8) were trained to self-administer a 0.8% w/v sucrose solution (0.8S). To minimize animal numbers, rats in Experiment 3 were also used for Experiment 4. Following 3 weekly saline habituation injections, caffeine (0, 5, 10, 20 mg/kg) was administered 30 min prior to a sucrose self-administration session using a within-subjects, Latin Square design. Caffeine testing procedures matched those described in Experiment 1.

#### 2.4.6. Experiment 4. Effect of acute caffeine administration on motivation for sucrose

Four weeks following completion of caffeine testing (Experiment 3), the effect of caffeine on motivation for a 0.8S sucrose reinforcer was evaluated using a PR1 schedule of reinforcement. Subjects received once-weekly injections of caffeine (0, 5, or 10 mg/kg) 30 min prior to the PR session. Caffeine testing and PR procedures matched those described in Experiment 2.

#### 2.4.7. Experiment 5. Evaluating the contribution of dopamine D_2_ receptors to caffeine-induced increases in operant responding for a sweetened alcohol solution

In order to evaluate the contribution of dopamine D_2_Rs to caffeine-induced increases in operant responding for a 10A2S solution, the dopamine D_2_R antagonist eticlopride was administered in conjunction with caffeine. For these studies, a naïve cohort of alcohol-trained (10A2S) Long-Evans rats (n = 8) was first habituated to double saline injections, with rats receiving 3-4 habituation injections twice weekly over 2 weeks. Eticlopride (0, 5, 10, 50 μg/kg) and caffeine (0, 5 mg/kg) were administered as separate injections 30 min before an operant self-administration session, with eticlopride administered first. Eticlopride doses and pretreatment times were chosen based upon previous studies (Arolfo et al., 2004; Liu and Weiss, 2002). Based on the results of Experiments 1 and 2, a 5 mg/kg dose of caffeine was used as it was the lowest effective dose that increased alcohol-reinforced responding without eliciting an increase in general motor activity (as measured by inactive lever responses). A within-subjects design was used for this study, with dose order randomized using a Latin Square design. Testing occurred once per week.

#### 2.4.8. Experiment 6. Further evaluating the contribution of dopamine D_2_ receptors to caffeine-induced increases in the reinforcing efficacy of alcohol

To further explore the contribution of D_2_Rs to caffeine’s reinforcement-enhancing effects, a moderate dose of eticlopride (10 μg/kg) was evaluated for its effects on caffeine-induced increases in alcohol self-administration and motivation for a 10A2S solution. In order to minimize animal numbers, Long-Evans rats (n = 8) that were used in Experiments 1 and 2 were used in these studies. Behavioral testing began approximately 2.5 months after the completion of Experiment 2.

For this study, rats were habituated to double saline injections, with habituation injections occurring once weekly for 2 weeks. Eticlopride (0, 10 μg/kg) and caffeine (0, 5 mg/kg) were injected 30 min prior to the operant self-administration session (FR2) for 10A2S, with dose order randomized across subjects using a Latin Square design. Testing occurred once weekly. Three weeks following the completion of FR testing, rats were tested for the effect of eticlopride on caffeine-induced increases in motivation for 10A2S using a PR1 schedule of reinforcement. Eticlopride (0, 10 μg/kg) and caffeine (0, 5 mg/kg) were injected 30 min before a PR1 session, with dose order randomized across subjects. PR testing occurred no more than once per week.

### 2.5. Data analysis

Data were analyzed using SPSS statistical software (SPSS Statistics 25, IBM). For Experiments 1-4, the effect of caffeine on alcohol- or sucrose-reinforced responses, alcohol or sucrose intake, break point, and inactive lever responses were analyzed using a one-way repeated measures analysis of variance (RM-ANOVA). Bonferroni-corrected pairwise comparisons were used to further examine significant effects. In order to examine time-dependent effects of caffeine on alcohol- and sucrose-reinforced responding, cumulative responses over the 30 min session (broken down into 5-min time bins) were analyzed using a two-way, caffeine dose X time RM-ANOVA. Significant main effects were examined with Bonferroni-corrected pairwise comparisons, whereas Bonferroni-corrected simple main effects (SME) analyses were used to explore any significant interactions. For Experiments 5 and 6, a two-way, eticlopride dose X caffeine dose RM-ANOVA was used, whereas time-dependent changes in response rate were examined using a three-way eticlopride dose X caffeine dose X time RM-ANOVA. Significant main effects were examined using Bonferroni-corrected pairwise comparisons, and interactions were explored using Bonferroni-corrected SME analyses. For all RM-ANOVAs, Mauchly’s test was used to test the assumption of sphericity; any violations of sphericity were corrected using the Greenhouse-Geisser procedure. Comparisons of baseline responding (Experiments 1 and 3) and break points (Experiments 2 and 4) between rats trained to self-administer a 10A2S or 0.8S solution were examined using independent-samples t-tests. For all analyses, alpha was set *a priori* at α= .05.

## 3. Results

### 3.1. Experiment 1. Effect of acute caffeine administration on operant responding for a sweetened alcohol solution

Baseline operant responding (mean ± SEM) for 10A2S over the 2 sessions preceding the start of caffeine testing was 86.75 (± 14.45), with an average alcohol intake of 0.57 (± 0.09) g/kg. Equipment malfunction on the last day of caffeine testing required 4 rats to be re-tested 1 week later for their final dose of caffeine (1 each from the 0, 5, 10, and 20 mg/kg caffeine doses).

As shown in Fig. 1A-B, low to moderate doses of caffeine significantly increased operant responding for, and intake of, a sweetened alcohol solution (10A2S). Results of a RM-ANOVA revealed a significant main effect of caffeine on the total number of alcohol-reinforced responses (F3,21=6.01, p=.004), as well as on alcohol intake (g/kg; F_3,21_=5.52, p=.006). Both the 5 mg/kg (ps<.03) and 10 mg/kg (ps<.01) doses of caffeine increased total session responses for 10A2S and alcohol intake, relative to the vehicle control. Although there was a significant main effect of caffeine on the number of inactive lever responses over the 30-min session, pairwise comparisons revealed no significant increase in inactive lever responses after caffeine administration, relative to vehicle (see Table 1 for group means and statistics).

**Figure 1.**
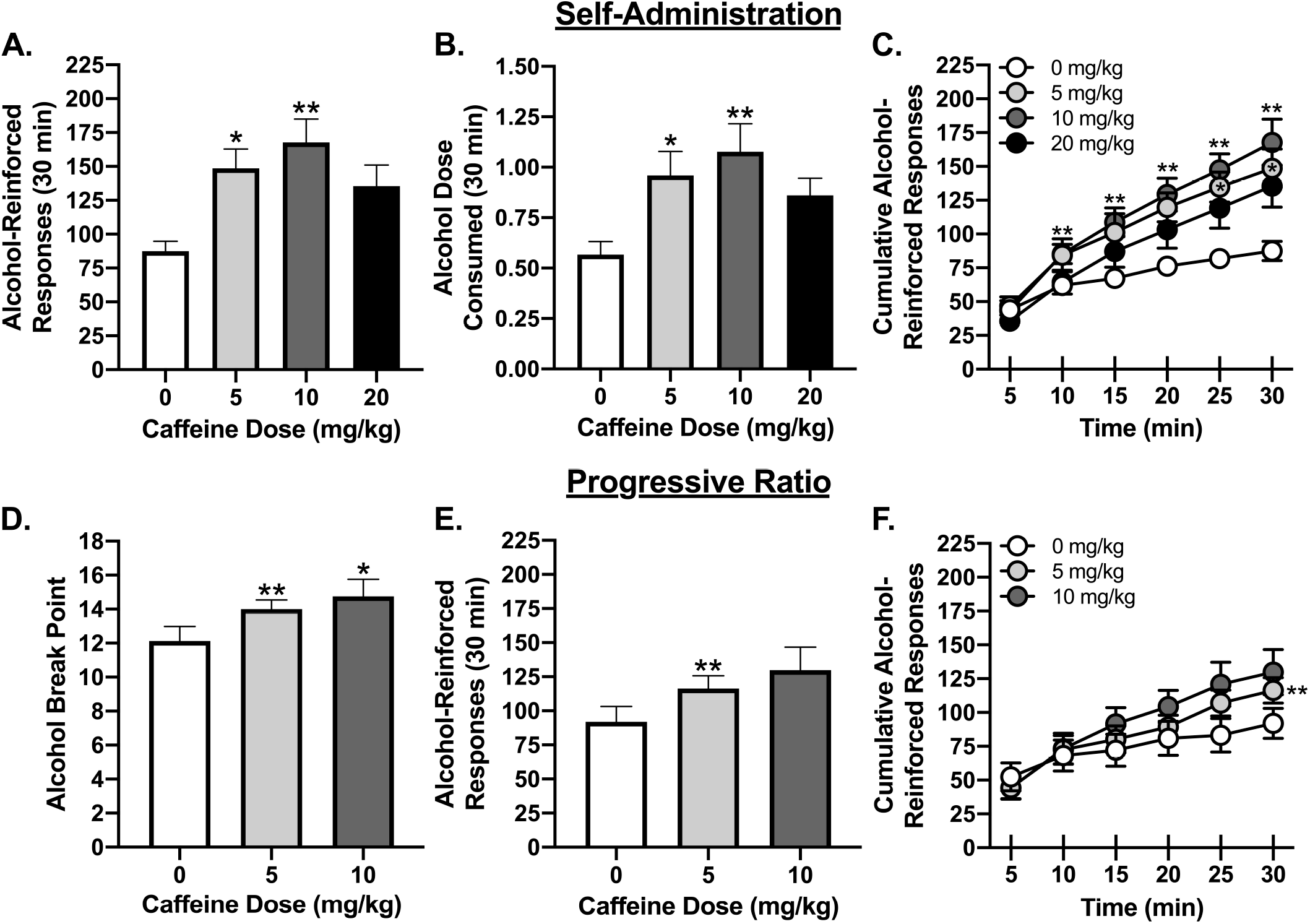
Caffeine increases operant responding and motivation for a sweetened alcohol reinforcer. (A) Total alcohol-reinforced responses and (B) alcohol dose consumed (g/kg) after caffeine administration, as measured on an FR2 schedule of reinforcement. (C) Cumulative responding for the sweetened alcohol reinforcer during the FR2 session, with responding broken down into 5-min time bins. (D) Alcohol break point and (E) total alcohol-reinforced responses following caffeine administration, as measured on a PR1 schedule of reinforcement. (F) Cumulative responding for sweetened alcohol during the PR session. Error bars represent standard error. Significant increase from vehicle control, *p < .05, **p < .01.

To determine whether caffeine altered the temporal pattern of responding for the sweetened alcohol reinforcer, cumulative responding over the 30-min session was evaluated (Fig. 1C). A caffeine dose X time RM-ANOVA revealed a significant main effect of caffeine dose (F_3,21_=4.89, p=.01) and time (F_1.3,8.9_=67.73, p<.001), as well as a significant caffeine dose X time interaction (F_15,105_=4.45, p<.001). The 10 mg/kg dose of caffeine increased the rate of alcohol-reinforced responding starting at 5 min and response rate remained elevated above vehicle for the remainder of the 30-min session (ps<.01). However, a 5 mg/kg dose of caffeine only significantly increased response rate during the last 10 min of the operant session (ps<.03).

### 3.2. Experiment 2. Effect of acute caffeine administration on motivation for a sweetened alcohol reinforcer

As hypothesized, caffeine likewise increased motivation for a 10A2S reinforcer (Fig. 1D-F). Results of a RM-ANOVA revealed a significant main effect of caffeine on alcohol break point (F_2,14_=7.85, p=.005), with both the 5 and 10 mg/kg caffeine doses increasing break point above vehicle control (ps<.05). There was also a significant main effect of caffeine on the total number of alcohol-reinforced responses over the 30-min PR session (F_1.2,8.2_=7.65, p=.02), with the 5 mg/kg caffeine dose significantly increasing total responses for 10A2S above vehicle control (p=.002). Although the 10 mg/kg caffeine dose also increased total responses for 10A2S, this effect was approaching significance (p=.054). There was no effect of caffeine on inactive lever responses (Table 1).

An analysis of cumulative response rate over time did not reveal a significant main effect of caffeine dose (F_2,14_=3.23, n.s.); however, there was a significant main effect of time (F_1.6,10.9_=47.89, p<.001), and a significant caffeine dose X time interaction (F_10,70_=5.43, p<.001). SME analyses revealed a significant caffeine-induced increase in response rate only during the last 5 min of the PR session, with a low dose of caffeine (5 mg/kg) significantly increasing response rate over vehicle control (p=.002). Despite a visible increase in response rate with the 10 mg/kg caffeine dose (see Fig. 1F), this effect was only approaching significance (p=.054).

### 3.3. Experiment 3. Effect of acute caffeine administration on operant responding for sucrose

Baseline responding for 0.8S over the 2 d immediately prior to the start of caffeine testing was 116.13 ± 19.07, with an average intake per session of 10.81 ± 1.97 ml/kg. There was no significant difference in baseline response rates between the 10A2S (Experiment 1) and 0.8S (Experiment 3) reinforcers, as revealed by an independent-samples t-test (t_14_=1.23, n.s.).

As shown in Fig. 2A-B, caffeine did not alter the total number of sucrose-reinforced responses (F_3,21_=0.80, n.s.) over the 30-min session. Caffeine likewise did not alter sucrose intake (F_3,21_=1.20, n.s.). A caffeine dose X time RM-ANOVA on cumulative response rate revealed a significant main effect of time (F_1.2,8.7_=80.97, p<.001), but as shown in Fig. 2C, there was no main effect of caffeine dose (F_3,21_=1.59, n.s.), and no caffeine dose X time interaction (F_15,105_=1.04, n.s). There was no effect of caffeine on inactive lever responses (Table 1).

**Figure 2.**
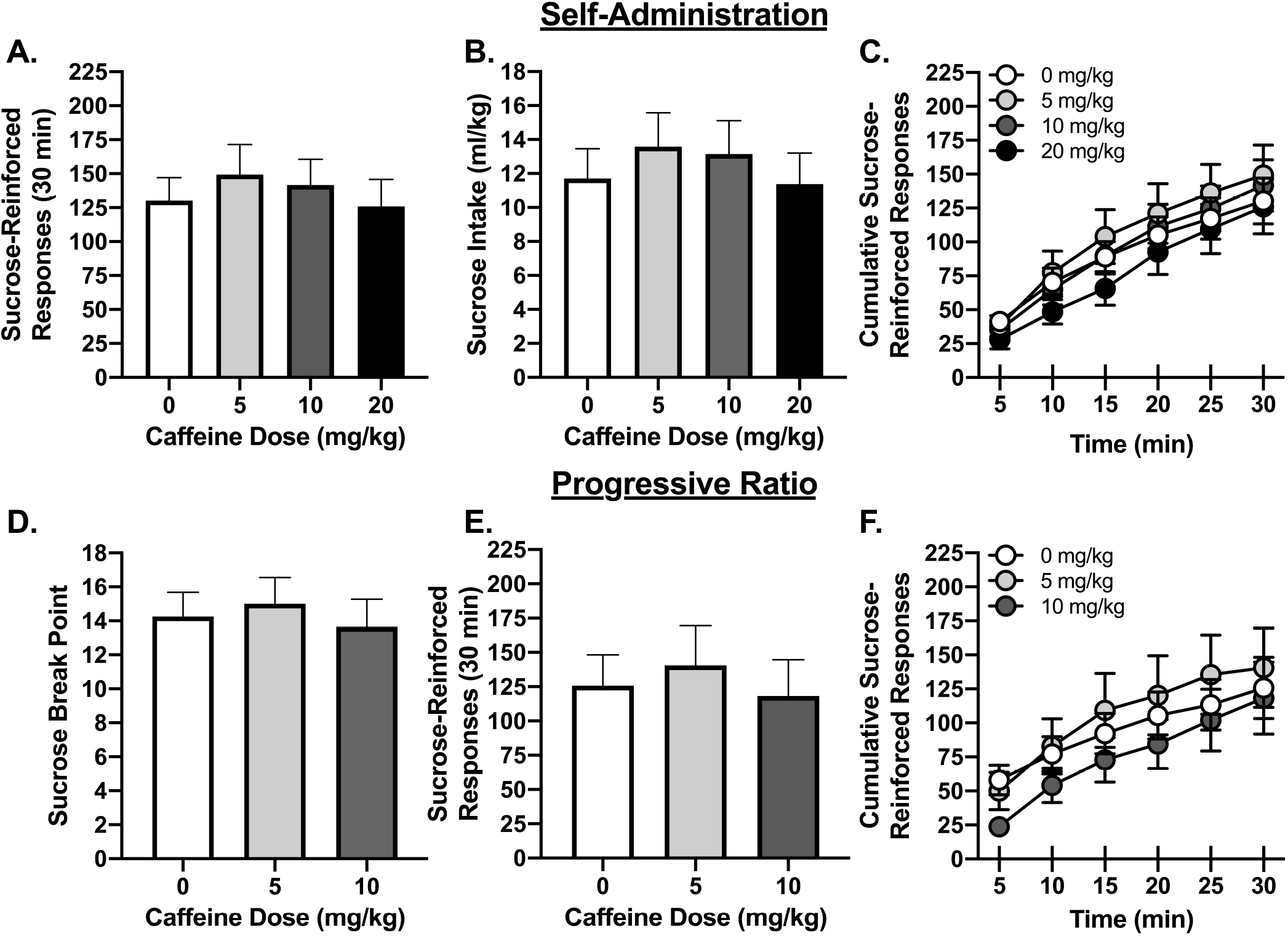
Caffeine does not alter sucrose-reinforced responding or motivation for sucrose. (A) Total sucrose-reinforced responses and (B) sucrose intake after caffeine administration, as measured on an FR2 schedule of reinforcement. (C) Cumulative responding for a sucrose reinforcer during the FR2 session, with responding broken down into 5-min time bins. (D) Sucrose break point and (E) total responses for sucrose following caffeine administration, as measured on a PR1 schedule of reinforcement. (F) Cumulative responding for sucrose during the PR session, with responding broken down into 5-min time bins. Error bars represent standard error.

### 3.4. Experiment 4. Effect of acute caffeine administration on motivation for sucrose

In contrast to the effect of caffeine on motivation for a sweetened alcohol reinforcer (Experiment 2), caffeine did not increase motivation for sucrose (see Fig. 2D-F). An independent-samples t-test confirmed that there was no significant difference in break point between the 10A2S (Experiment 1) and 0.8S (Experiment 3) reinforcers (t_11.4_=1.27, n.s.), suggesting that the relative reinforcing efficacy of the two solutions was equivalent. Results of a RM-ANOVA revealed no significant effect of caffeine on sucrose break point (F_2,14_=1.08, n.s.) or on the total number of sucrose-reinforced responses (F_2,14_=0.87, n.s.). Likewise, caffeine did not alter the overall rate of sucrose-reinforced responding across the 30-min PR session. A caffeine dose X time RM-ANOVA revealed no significant main effect of caffeine dose (F_2,14_=2.38, n.s.) on cumulative response rate; however, there was a significant main effect of time (F_1.1,8.0_=22.90, p<.001), as well as a significant caffeine dose X time interaction (F_10,70_=2.08, p=.04). However, SME analyses revealed only a marginal, but non-significant, effect of caffeine during the first 5 min (p=.054), with no significant effect of caffeine dose on response rate at any other time point. Caffeine did not alter inactive lever responses (Table 1).

### 3.5. Experiment 5. Evaluating the contribution of dopamine D_2_ receptors to caffeine-induced increases in operant responding for a sweetened alcohol solution

Responding (mean ± SEM) over the 2 baseline self-administration sessions immediately prior to the start of eticlopride and caffeine testing was 101.94 (± 7.13), with an average alcohol intake of 0.74 (± 0.04) g/kg. Injection problems mid-way through the experiment required 2 rats to be re-tested at the end of the study (1 each from 0 μg/kg eticlopride + 5 mg/kg caffeine and 5 μg/kg eticlopride + 5 mg/kg caffeine).

Consistent with Experiment 1, caffeine (5 mg/kg) significantly increased alcohol-reinforced responding in rats. However, despite the fact that eticlopride reduced operant responding for a 10A2S reinforcer on its own, eticlopride was unable to reverse a caffeine-induced increase in alcohol-reinforced responding. This finding was confirmed by an eticlopride dose X caffeine dose RM-ANOVA, which showed a significant main effect of eticlopride dose (F_3,21_=8.27, p=.001) and caffeine dose (F_1,7_=56.24, p<.001) on the total number of alcohol-reinforced responses. As shown in Fig. 3A, caffeine (5 mg/kg) increased total session responses for 10A2S, whereas 50 μg/kg eticlopride decreased total session responses below the vehicle control (p=.003). However, there was no significant eticlopride dose X caffeine dose interaction (F_3,21_=0.91, n.s.), demonstrating that eticlopride was not able to reverse a caffeine-induced increase in alcohol-reinforced responding in rats. A similar pattern of results was observed for alcohol intake, with significant main effects of eticlopride dose (F_3,21_=10.22, p<.001) and caffeine dose (F_1,7_=65.13, p<.001), but no significant eticlopride dose X caffeine dose interaction (F_3,21_=0.99, n.s.). As shown in Fig. 3B, although caffeine (5 mg/kg) increased alcohol intake, and eticlopride (50 μg/kg) significantly reduced alcohol intake, relative to the vehicle control (p=.003), eticlopride did not block a caffeine-induced increase in alcohol consumption. Caffeine did elicit a small increase in inactive lever responses (see Table 2); however, there was no significant effect of eticlopride, nor a significant eticlopride dose X caffeine dose interaction.

**Figure 3.**
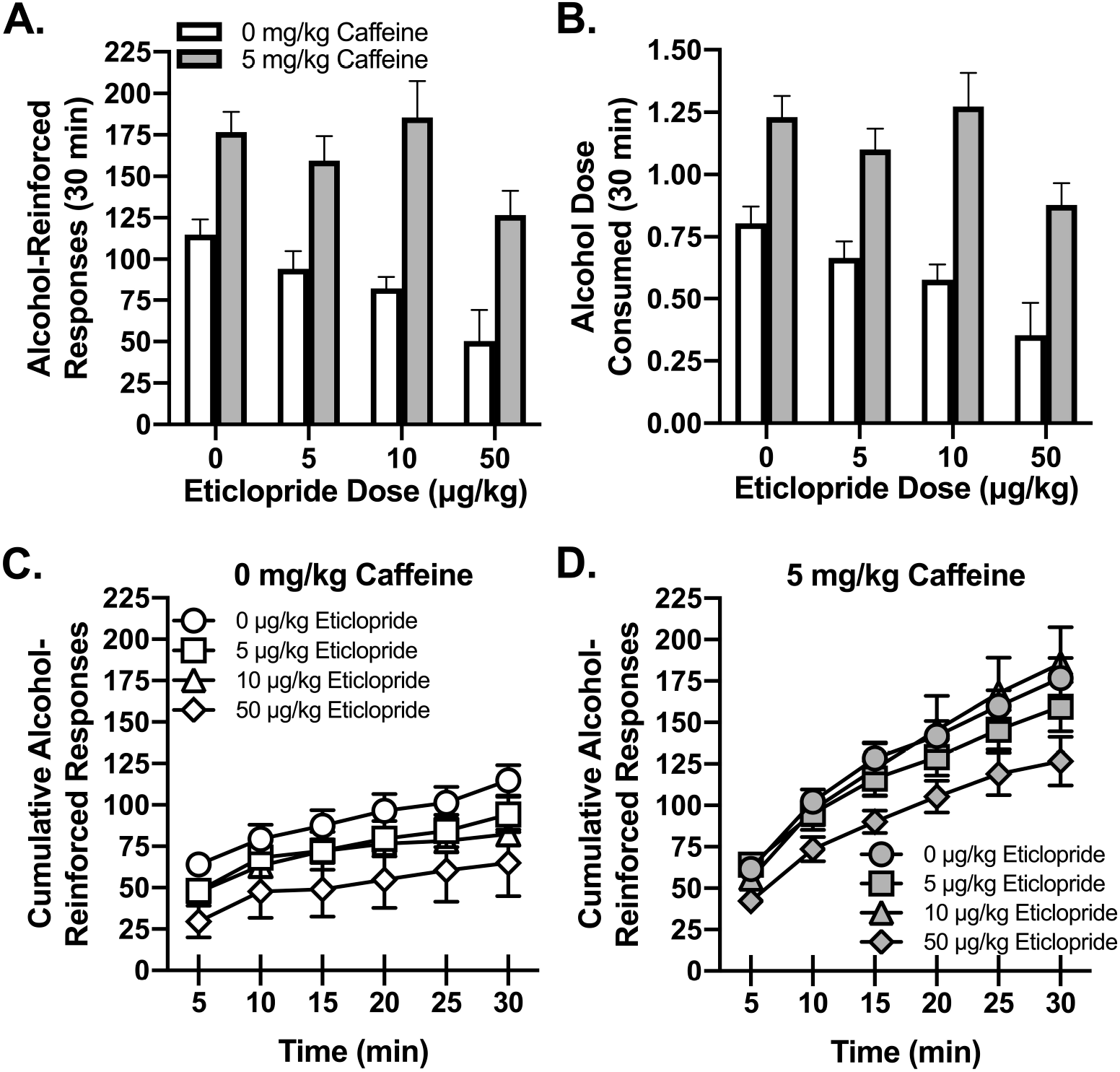
Eticlopride does not inhibit a caffeine-induced increase in alcohol self-administration. (A) Total alcohol-reinforced responses and (B) alcohol dose consumed (g/kg) following eticlopride and caffeine administration. Effect of eticlopride on the rate of responding for 10A2S in (C) vehicle-treated rats (0 mg/kg caffeine) and (D) caffeine-treated rats (5 mg/kg). Cumulative response rate was broken down into 5-min time bins. Error bars represent standard error.

To determine whether eticlopride, caffeine, or their combination, altered the temporal pattern of responding for 10A2S, an eticlopride dose X caffeine dose X time RM-ANOVA was conducted. Analyses revealed a significant main effect of eticlopride dose (F_3,21_=4.59, p=.01), caffeine dose (F_1,7_=38.07, p<.001), and time (F_2.3,16.3_=104.44, p<.001), as well as a significant caffeine dose X time interaction (F_2.2,15.2_=42.13, p<.001). Caffeine (5 mg/kg) elicited an increase in alcohol-reinforced response rate that emerged after 5 min and remained elevated for the remainder of the session (ps<.02). However, there was no significant interactions of eticlopride dose and time (F_15,105_=1.53, n.s.), eticlopride dose and caffeine dose (F_1.6,11.3_=0.62, n.s.), or eticlopride dose, caffeine dose, and time (F_15,105_=1.33, n.s.).

### 3.6. Experiment 6. Further evaluating the contribution of dopamine D_2_ receptors to caffeine-induced increases in the reinforcing efficacy of alcohol

To further examine the contribution of D_2_Rs to the reinforcement-enhancing effects of caffeine, the effect of a moderate dose of eticlopride (10 μg/kg), in combination with caffeine, on alcohol-reinforced responding was examined using both an FR and a PR schedule of reinforcement. As shown in Fig. 4A-B, we were able to replicate the results of Experiment 5 demonstrating that eticlopride was unable to reverse a caffeine-induced increase in alcohol self-administration using an FR2 schedule of reinforcement. Results of an eticlopride dose X caffeine dose RM-ANOVA confirmed a significant main effect of eticlopride dose (F_1,7_=10.47, p=.01) and caffeine dose (F_1,7_=16.00, p=.005) on alcohol-reinforced responding, but no significant eticlopride dose X caffeine dose interaction (F_1,7_=0.08, n.s.). Similar results were observed for alcohol intake, with a significant main effect of eticlopride dose (F_1,7_=9.70, p=.02) and caffeine dose (F_1,7_=16.62, p=.005), but no significant eticlopride dose X caffeine dose interaction (F_1,7_=0.04, n.s.). Eticlopride did not alter a caffeine-induced increase in response rate over the 30-min session (data not shown).

**Figure 4.**
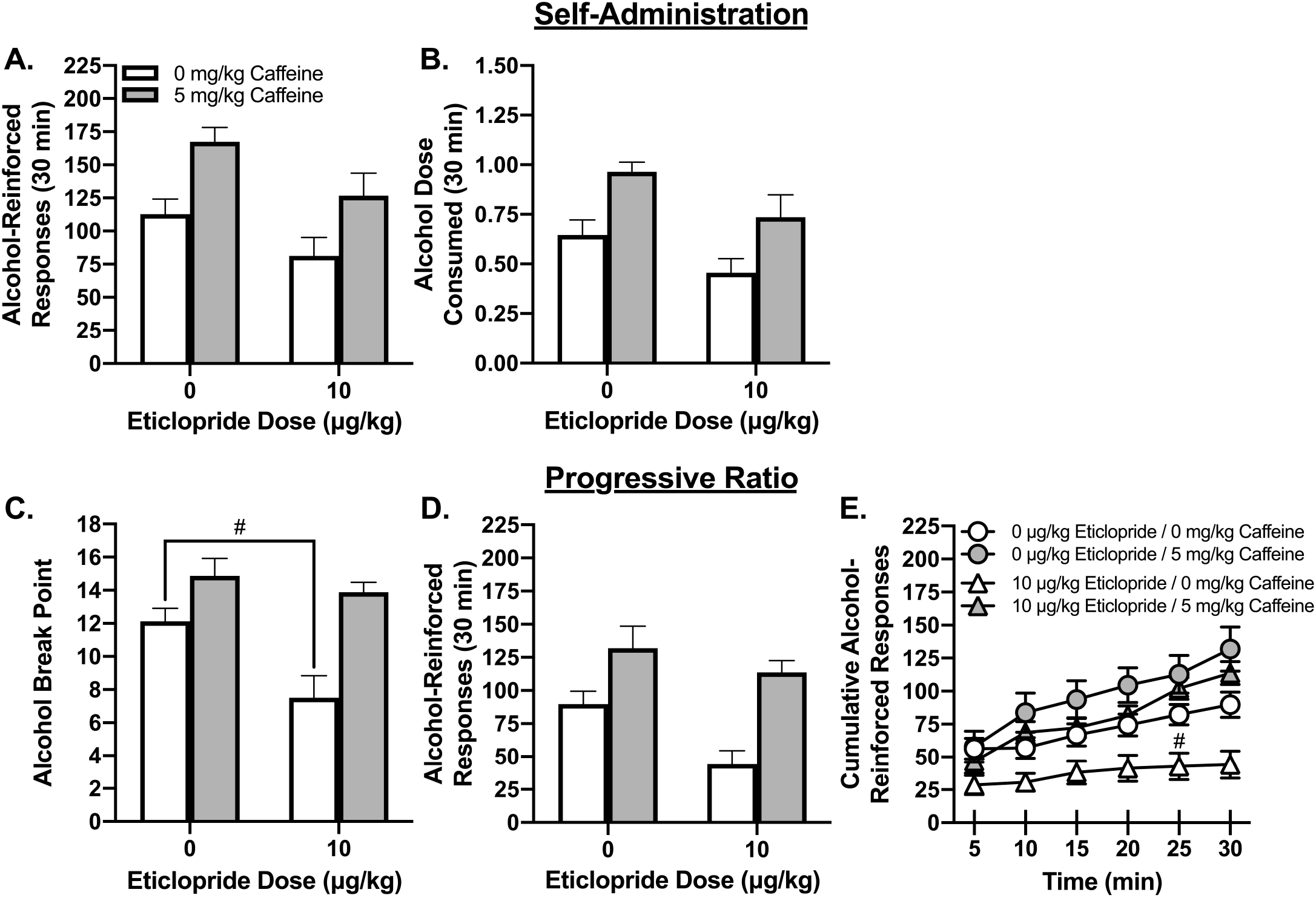
Eticlopride does not attenuate the reinforcement-enhancing effects of caffeine. (A) Total alcohol-reinforced responses and (B) alcohol dose consumed (g/kg) following eticlopride and caffeine administration, as measured on an FR2 schedule of reinforcement. (C) Alcohol break point and (D) alcohol-reinforced responses following eticlopride and caffeine administration, as measured on a PR1 schedule of reinforcement. (E) Cumulative alcohol-reinforced responses during the PR1 session, with responding broken down into 5-min time bins. Error bars represent standard error. # Significant difference between the 0 μg/kg eticlopride + 0 mg/kg caffeine and the 10 μg/kg eticlopride + 0 mg/kg caffeine groups, p < .05.

To address the impact of eticlopride on caffeine-induced increases in motivation for a 10A2S reinforcer, a PR1 schedule of reinforcement was used. Consistent with the self-administration data, eticlopride did not block a caffeine-induced increase in motivation for a 10A2S reinforcer (see Fig. 4C-E). An eticlopride dose X caffeine dose RM-ANOVA revealed a significant main effect of eticlopride dose (F_1,7_=5.68, p=.049) and caffeine dose (F_1,7_=34.57, p=.001) on break point, as well as a significant eticlopride dose X caffeine dose interaction (F_1,7_=9.70, p=.02). However, SME analyses revealed that eticlopride reduced break point in rats administered 0 mg/kg caffeine (p=.02), but did not alter break point in rats administered 5 mg/kg caffeine (p=.34). A similar pattern of results was obtained for the total number of alcohol-reinforced responses, with significant main effects of eticlopride dose (F_1,7_=5.86, p=.046) and caffeine dose (F_1,7_=38.28, p<.001). However, there was no significant eticlopride dose X caffeine dose interaction for total session alcohol responses (F_1,7_=4.4, n.s.). There was no significant effect of eticlopride, caffeine, or their combination, on inactive lever responses (Table 2).

In an analysis of response rate across the 30-min PR session, an eticlopride dose X caffeine dose X time RM-ANOVA revealed a significant main effect of eticlopride dose (F_1,7_=9.89, p=.02), caffeine dose (F_1,7_=29.81, p=.001), and time (F_1.3,8.8_=42.25, p<.001), as well as a significant caffeine dose X time interaction (F_1.9,13.1_=16.86, p<.001), with a caffeine-induced increase in response rate emerging after 5 min (ps<.01). Although there were no significant interactions of eticlopride dose and time (F_1.5,10.7_=0.68, n.s.) or eticlopride dose and caffeine dose (F_1,7_=3.92, n.s.), there was a significant eticlopride dose X caffeine dose X time interaction (F_5,35_=2.55, p=.046). However, SME analyses revealed that a significant eticlopride dose X caffeine dose interaction was restricted to the 25-min time point (p=.04), with eticlopride reducing response rate in vehicle-treated rats (0 mg/kg caffeine; p=.01), but not in caffeine-treated rats (5 mg/kg caffeine; p=.47).

## 4. Discussion

In the current study, we examined the hypothesis that caffeine may be increasing alcohol intake by enhancing the reinforcing efficacy of alcohol using an operant conditioning paradigm. As hypothesized, low to moderate doses of caffeine (5-10 mg/kg) significantly increased operant alcohol self-administration and motivation for a sweetened alcohol reinforcer. These effects appear to be selective for alcohol, as caffeine did not likewise increase operant responding for, or motivation for, a behaviorally-matched concentration of sucrose. However, in contrast to previous hypotheses explaining caffeine’s reinforcement-enhancing effects (Ferré, 2016, 2013; Ferré and O’Brien, 2011), systemic blockade of dopamine D_2_Rs did not inhibit a caffeine-induced increase in operant responding for a sweetened alcohol reinforcer. These results suggest that caffeine is increasing alcohol intake, in part, by enhancing the reinforcing efficacy of alcohol, an effect which may be independent of increased activity at the dopamine D_2_R.

### 4.1. Caffeine increases the reinforcing efficacy of alcohol

Although caffeine has previously been reported to increase alcohol intake in rodent models (Franklin et al., 2013; Fritz et al., 2016; Kunin et al., 2000; Okhuarobo et al., 2018; Rezvani et al., 2013; SanMiguel et al., 2019), the current study is among the first to show a direct effect of caffeine on the positive reinforcing effects of alcohol. As shown in Experiment 1, caffeine increased operant responding for a sweetened alcohol reinforcer, as well as alcohol intake, an effect which was only observed after low to moderate doses of caffeine. However, this increase in alcohol-reinforced responding emerged after 5 min, at timepoints when vehicle-treated rats began to show a deceleration in alcohol intake. These results suggest that caffeine is not altering initial alcohol-seeking behaviors, but rather alters consummatory processes by influencing the maintenance of alcohol self-administration and/or inhibiting satiety-related processes that serve to limit alcohol intake (Samson and Czachowski, 2003; Samson and Hodge, 1996). Alcohol-induced increases in extracellular adenosine concentrations have been proposed to impose a “brake” on the reinforcing effects of alcohol via an inhibition of dopamine signaling (Ferré et al., 2018a; Ferré and O’Brien, 2011; Sharma et al., 2010). By selectively altering consummatory-related behaviors, caffeine and other adenosine receptor antagonists may remove this “brake”, leading to a loss of control over drinking behavior and potentially dangerous levels of alcohol intake (Ferré, 2016; Marczinski, 2014).

Changes in FR responding can be conceptually difficult to interpret, however, as caffeine-induced increases in operant responding could reflect an increase in the reinforcing efficacy of alcohol or an inhibition of alcohol’s pharmacological effects (Faccidomo et al., 2009). PR schedules can help to further interpret changes in operant responding by evaluating motivation, or the level of effort a subject is willing to put in for alcohol (Richardson and Roberts, 1996). Consistent with our hypothesis, we found that low to moderate doses of caffeine elicited a small but significant increase in alcohol break point, supporting a caffeine-induced increase in motivation for a sweetened alcohol reinforcer. Since subjects were responding at higher rates for, and exerting additional effort toward, gaining a sweetened alcohol reinforcer, it can be inferred that the reinforcing efficacy of alcohol was enhanced by caffeine (Bickel et al., 2000; Nehlig, 1999; Stafford et al., 1998). As such, caffeine may be enhancing the reinforcement or pleasure experienced with alcohol, leading to increased (and prolonged) alcohol intake and enhanced motivation for alcohol. Importantly, these results are consistent with previous studies reporting an increase in operant responding for alcohol when combined with a caffeinated ED (Roldán et al., 2018). However, EDs combine caffeine along with other potentially psychoactive ingredients (De Sanctis et al., 2017; Seifert et al., 2011). The current study adds to this literature by demonstrating an immediate and direct pharmacological action of caffeine on the positive reinforcing effects of alcohol.

Interestingly, these reinforcement-enhancing effects of caffeine appear to be specific for alcohol. For the current series of studies, a low concentration of sucrose (2%) was included in the alcohol solution (Experiments 1-2 and 5-6) to encourage moderate and pharmacologically-relevant levels of alcohol intake (0.5-0.8 g/kg). However, this created a potential confound in our interpretation of the alcohol self-administration data and it was unclear whether caffeine was increasing the reinforcing efficacy of alcohol or sucrose. Therefore, in Experiments 3 and 4, responding for a concentration of sucrose (0.8%) that elicited a similar level of responding to, and motivation for, a 10A2S reinforcer (and therefore was putatively equivalent in its reinforcing efficacy to a 10A2S solution) was tested. Contrary to the previous literature (Retzbach et al., 2014; Sheppard et al., 2012), caffeine did not increase sucrose-reinforced responding in rats, nor did caffeine increase sucrose break point. These results were surprising, as caffeine has been proposed to act as a reinforcement enhancer (Garber and Lustig, 2011; Prieto et al., 2016; Sheppard et al., 2012), and has been reported to increase the reinforcing efficacy of both drug (Gannon et al., 2019, 2018; Prieto et al., 2016; Rezvani et al., 2013; Schenk et al., 1994; Shoaib et al., 1999) and non-drug (Bradley and Palmatier, 2019; Retzbach et al., 2014; Sheppard et al., 2012) reinforcers. However, caffeine-induced increases in the reinforcing efficacy of sucrose were reported for higher sucrose concentrations (Sheppard et al., 2012) and/or following repeated caffeine administration (Retzbach et al., 2014). The current results do not necessarily negate the hypothesis that caffeine acts as a generalized reinforcement enhancer, but rather might suggest that caffeine’s reinforcement-enhancing effects are more immediate and more pronounced for a drug reinforcer, such as alcohol.

Although these results can be interpreted to mean that caffeine increases alcohol intake, in part, by increasing the reinforcing efficacy of alcohol, it is important to rule out potential alternative explanations. Caffeine, for instance, is a modest psychostimulant and elicits increases in motor activity at moderate doses (10-30 mg/kg) in rats (Garrett and Holtzman, 1994; Powell et al., 2001). Although it could be argued that caffeine-induced increases in operant responding for alcohol simply reflect a nonspecific motor stimulant response to caffeine, we believe this explanation is unlikely. First, caffeine had little to no effect on inactive lever responses, and second, caffeine’s reinforcement enhancing effects were selective for alcohol, as caffeine did not likewise increase operant responding for a sucrose reinforcer. Finally, D_2_R antagonists, including eticlopride, attenuate the motor stimulant effects of caffeine (Garrett and Holtzman, 1994; Powell et al., 2001). Yet, in Experiments 5 and 6, caffeine was still able to elicit a significant increase in alcohol-reinforced responding, even at doses that were found to inhibit the stimulant actions of caffeine (Garrett and Holtzman, 1994).

Alternatively, researchers have hypothesized that caffeinated EDs increase alcohol intake by attenuating the intoxicating and sedative-hypnotic effects of alcohol (Ferré, 2016; Ferré and O’Brien, 2011; Marczinski, 2011). In line with this hypothesis, caffeine attenuates the motor depressant (Ferreira et al., 2004; Fritz et al., 2014), motor incoordinating (Connole et al., 2004; Dar, 1988; Dar et al., 1987; Fritz et al., 2014), and somnogenic effects of alcohol (Dar et al., 1987; El Yacoubi, 2003; Fang et al., 2017). However, little direct evidence supports this hypothesis in humans (see Benson et al., 2014; Johnson et al., 2018) and we do not believe this is a likely explanation for the increase in alcohol-reinforced responding observed in the current study. First, caffeine-induced increases in alcohol-reinforced responding emerged after only 5 min – a time when brain alcohol levels would still be rising (Doyon et al., 2005). It is unlikely that subjects would be experiencing a sedative response to consumed alcohol at this early time point, as previous studies demonstrated that a motor depressant response to consumed alcohol only emerged after 20 min (Fritz et al., 2014). Second, caffeine elicited an increase in motivation for alcohol using a PR schedule of reinforcement despite the low level of alcohol intake observed in these studies (mean ± SEM alcohol intake was 0.16 ± 0.01 g/kg for vehicle-treated rats in Experiment 2). If caffeine was increasing alcohol intake simply by masking the intoxicating and/or somnogenic effects of alcohol, we would not expect to see an increase in operant responding under a PR schedule of reinforcement given the low, non-sedative doses of alcohol consumed in these types of studies.

### 4.2. Evaluating the contribution of dopamine D_2_ receptors to caffeine’s reinforcement-enhancing effects

In order to examine the neuropharmacological mechanisms underlying caffeine’s reinforcement-enhancing effects, we investigated a potential role for dopamine D_2_Rs. Although D_2_R function is critical for the maintenance of alcohol self-administration (Arolfo et al., 2004; Files, 1998; Hodge et al., 1997), it has also been proposed to contribute to the reinforcement-enhancing effects of caffeine (Ferré, 2013, 2016; Ferré and O’Brien, 2011; Green and Schenk, 2002). Within the striatum, A_2A_Rs are predominantly coupled with D_2_Rs as an A_2A_R-D_2_R heterotetrameric complex (Ferré et al., 2018b, 2018a). Caffeine and other A_2A_R antagonists increase the affinity and efficacy of dopamine at the D_2_R, thereby putatively enhancing the response to rewarding stimuli, including alcohol (Bonaventura et al., 2015; Ferré et al., 2018b, 2018a; Volkow et al., 2015). We therefore hypothesized that a caffeine-induced increase in D_2_R function could be mediating the reinforcement-enhancing effects of caffeine.

In Experiments 5 and 6 we used the D_2_R antagonist eticlopride to test whether an inhibition of D_2_R function could reverse the behavioral effects of caffeine and prevent a caffeine-induced increase in operant responding for alcohol. Consistent with previous studies (Arolfo et al., 2004), eticlopride decreased alcohol-reinforced responding and alcohol intake, as well as alcohol break point, confirming a critical role of D_2_Rs to the positive reinforcing effects of alcohol (Arolfo et al., 2004; Czachowski et al., 2001; Files, 1998; Hodge et al., 1997; Risinger et al., 2000). Yet despite this inhibition of D_2_R function, caffeine continued to significantly increase both alcohol-reinforced responding (Experiments 5 and 6) and alcohol break point (Experiment 6) in rats. Importantly, this effect was observed even at a sedative dose of eticlopride (50 μg/kg; see Ferrari and Giuliani, 1995), demonstrating that caffeine can still increase the positive reinforcing effects of alcohol even as non-specific motor sedative effects of eticlopride emerge. Although these results lend support to the hypothesis that caffeine counteracts the sedative effects of eticlopride (Collins et al., 2010), they suggest that D_2_R function is not necessary for caffeine-induced increases in the reinforcing efficacy of alcohol.

Although caffeine-induced increases in the reinforcing efficacy of alcohol may occur independently of D_2_Rs, this does not necessarily negate a role of adenosine-dopamine interactions in caffeine’s reinforcement-enhancing effects. First, eticlopride was administered systemically in the current studies; a more specific contribution of D_2_Rs to caffeine’s reinforcement-enhancing effects may be observed if site-specific injections of eticlopride were used. Likewise, A_2A_Rs functionally interact with several different receptors, including D3, D4, CBi, and mGlu_5_ receptors (Wydra et al., 2020), suggesting other possible mechanistic explanations for the results observed in the current study. The A_2A_R-mGluR_5_ interaction is particularly intriguing, as mGluR_5_ has been found to regulate the positive reinforcing actions of alcohol (Besheer et al., 2008; Schroeder et al., 2005). Likewise, an mGluR_5_ antagonist, in combination with an A_2A_R antagonist, has been reported to regulate alcohol self-administration and seeking behaviors in rats (Adams et al., 2008). However, more work is needed to determine whether A_2A_R-mGluR_5_ heteromers (or other A_2A_R interactions) may better explain a caffeine-induced increase in alcohol self-administration. Caffeine is also an AiR antagonist (Daly and Fredholm, 1998; Fredholm et al., 1999), which can likewise form heteromeric complexes not only with A_2A_Rs, but also with DiRs (Ginés et al., 2000; Wydra et al., 2020). Although previous studies have highlighted a preferential role of A_2A_Rs in alcohol drinking and operant self-administration (Arolfo et al., 2004; SanMiguel et al., 2019), others have reported a preferential role of AiRs in alcohol drinking (Fritz and Boehm, 2015). As DiR antagonists likewise decrease self-administration (Hodge et al., 1997), a caffeine mediated alteration of AiR-DiR function may be a more likely candidate for caffeine’s reinforcement-enhancing effects.

### 4.3. Conclusion

In conclusion, the results of the current study demonstrate that caffeine selectively increases both the positive reinforcing effects of self-administered alcohol, as well as motivation for an alcohol reinforcer. Although the neuropharmacological mechanism underlying this reinforcement-enhancing effect remains to be discovered, these results further emphasize public health concerns regarding the combination of alcohol with caffeinated EDs (De Sanctis et al., 2017; Marczinski and Fillmore, 2014; Vida and Rácz, 2015). By promoting both motivation for alcohol as well as consummatory-related behaviors, caffeine may lead to a loss of control over alcohol intake, leading to hazardous drinking and increased risk for adverse consequences of heavy alcohol use.

## Acknowledgements

The authors would like to acknowledge Abigail Bracken, Krista Brady, Angela Cardillo, Emily Frantz, Logan Gregory, Erik Homberger, Megan McVeety, Ashmita Mukherjee, Kelly Rogawski, Elle Sarracco, Morgan Valle, and Lydia Yorks for their technical assistance on this project, as well as Dr. Joyce Besheer for her thoughtful comments and suggestions.

